# Intracellular Delivery of Bone Nanoparticles to Mitigate Irradiation-Induced Damage in Bone Marrow Mesenchymal Stem Cells

**DOI:** 10.1101/2025.05.05.652269

**Authors:** Bo Wang, Xiaolong Ma, Christopher Nguyen, Austin Stellpflug, Siqi Li, Lu Han, Anne Frei, Heather Himburg, Linxia Gu, Pengfei Dong, Shue Wang, Michael Mak, Rongxue Wu

## Abstract

Ionizing radiation (IR)-induced bone damage presents a major clinical challenge by impairing bone marrow function and disrupting normal bone remodeling. Bone regeneration depends on bone marrow-derived mesenchymal stem cells (BMSCs), which are highly sensitive to IR that causes DNA damage, oxidative stress, apoptosis, and a shift from osteogenesis to adipogenesis, ultimately leading to bone loss and impaired healing. This study evaluated the therapeutic potential of intracellularly delivered bone-derived nanoparticles (BPs) in mitigating IR-induced BMSCs damage. We found that IR exposure caused significant BMSCs dysfunction, including reduced proliferation, increased apoptosis, persistent DNA damage, and a shift toward adipogenic differentiation. Treatment with BPs led to efficient intracellular uptake, improved cell morphology, enhanced proliferation, reduced apoptosis, and preservation of balanced differentiation capacity. Transcriptomic analysis via RNA sequencing revealed that BPs restored key molecular pathways disrupted by IR, particularly those involved in cell cycle regulation, extracellular matrix (ECM) remodeling, and apoptosis. By reversing these transcriptional impairments, BPs supported genomic stability and the regenerative function of BMSCs. Overall, these findings suggest that BPs effectively counteract IR-induced cellular damage and enhance the regenerative capacity of BMSCs, offering a promising therapeutic strategy for radiation -induced skeletal injuries.

## 1. Introduction

Ionizing radiation (IR) originates from various sources, including nuclear accidents, radiotherapy, nuclear weapons, industrial processes, and space exploration. Prolonged or high-dose exposure to IR can severely damage cells by directly disrupting deoxyribonucleic acid (DNA) strands, leading to mutations and functional impairments. Additionally, IR generates reactive free radicals that exacerbate tissue damage, induce cell death, and trigger inflammatory responses. These combined effects disrupt tissue repair mechanisms, increasing the risk of chronic conditions such as fibrosis and cancer [1-4].

Bone tissue, due to its high calcium content, absorbs approximately 40% more IR than soft tissues, making it particularly vulnerable to IR-induced damage [5-7]. IR disrupts bone remodeling by inhibiting osteoblast activity while enhancing osteoclast function, leading to excessive bone resorption [8-12]. This imbalance weakens the bone matrix by impairing collagen synthesis and mineralization, compromising structural integrity. Furthermore, IR reduces vascularization in bone tissue, limiting blood flow and oxygen delivery necessary for repair and regeneration [13]. These effects contribute to progressive bone loss, decreased mechanical strength, and an increased risk of fractures [7, 14-16]. Acute effects include inflammation and refractory pain, while long-term complications such as delayed healing, nonunion, and osteoradionecrosis further hinder bone recovery and function [17-20].

Current therapeutic strategies for IR-induced bone damage primarily focus on symptom relief and structural support through interventions such as wound management, hyperbaric oxygen therapy (HBOT), surgical fixation, and bone grafting [21]. While these treatments can temporarily alleviate pain, control infections, provide mechanical stability, and repair bone defect, they offer limited potential for true bone regeneration and restoration of long-term bone quality [22, 23]. These limitations highlight the urgent need for innovative therapies that address the underlying biological disruptions caused by IR, promoting bone regeneration and functional recovery [24-27].

Bone regeneration is a complex process regulated by cellular interactions and signaling pathways, in which the bone marrow-derived mesenchymal stem cells (BMSCs) playing a key role by differentiating into osteoblasts and chondrocytes [28-31]. However, IR not only directly damages bone tissue but also disrupts the bone marrow microenvironment, affecting both hematopoietic stem cells (HSCs) and BMSCs in bone marrow [32-34]. IR exposure induces DNA damage, increases reactive oxygen species (ROS), promotes apoptosis, and accelerates BMSC senescence, shifting their differentiation from osteogenesis to adipogenesis, leading to bone loss and impaired healing [35]. Additionally, radiation-induced bone marrow damage triggers inflammation and alters critical signaling pathways, further exacerbating bone deterioration [36, 37]. Therefore, developing a targeted cellular therapy capable of restoring IR-induced BMSC damage and dysfunction is critical for preserving bone integrity and enhancing repair after radiation exposure.

Currently, there are no effective therapies exist to fully reverse IR-induced damage in BMSCs or restore their osteogenic regenerative potential. Conventional treatments, such as growth factor supplementation, pharmacological agents, and exogenous stem cell transplantation, are limited by poor bioavailability, immune rejection, and inconsistent therapeutic outcomes. Moreover, these approaches fail to directly address the intracellular signaling disruptions and oxidative stress that contribute to BMSC dysfunction after IR exposure [38- 40].

Recent advances in nanotechnology, particularly the use of nanoparticles (NPs), offer a promising strategy for enhancing stem cell therapies by enabling efficient intracellular uptake and targeted delivery of bioactive molecules. Their nanoscale size allows them to interact closely with cells, modulate the microenvironment, and mimic key biological functions that support regeneration. By facilitating controlled signaling, promoting cell survival, and guiding lineage-specific differentiation, NPs offer a powerful tool for precisely regulating stem cell fate and improving therapeutic outcomes in regenerative medicine [41-44].

Building on recent advancements in nanotechnology, our lab has developed a novel class of bone -derived nanoparticles (BPs) from decellularized porcine bone [45]. These BPs retain essential extracellular matrix (ECM) protein components of bone and possess nanoscale properties that enable efficient internalization by BMSCs. Once internalized, BPs enhance cell proliferation and significantly promote osteogenic differentiation under osteogenic conditions. Additionally, BPs exhibit strong potential as a grafting material, providing essential biochemical cues that stimulate cellular activity and support tissue remodeling [45, 46]. These properties position BPs as a powerful tool for bone tissue engineering, with the potential to address critical challenges in IR-induced bone damage.

Therefore, this study aims to evaluate the therapeutic potential of intracellularly delivered BPs in mitigating IR-induced damage in BMSCs. Specifically, we will assess whether BPs can restore BMSC function by enhancing cell survival, promoting proliferation, and stimulating osteogenic differentiation following IR exposure. Additionally, we will analyze gene expression patterns to identify the molecular pathways involved in BPs’ regenerative effects. The findings from this study could contribute to the development of NP-based cell therapies for enhancing bone repair and regeneration following IR exposure.

## 2. Materials and Methods

### 2.1. Bone decellularization and BP fabrication

Fresh porcine tibias (sourced from Medical College of Wisconsin -approved vendors) were processed following our established protocols [46-48]. Bones were sectioned into small pieces, fully demineralized with 0.5 N HCl (Sigma-Aldrich), and decellularized using a solution of 0.5% SDS and 1% Triton X-100 (Sigma-Aldrich). After thorough washing to remove residual detergents, the bone was lyophilized and milled into fine powders using a Thomas Wiley® Mini Cutting Mill (Thomas Scientific). The bone powders were then digested with 15% pepsin (Sigma-Aldrich) at pH 2.5 and 45°C for 2–3 days to create a bone stock solution with a protein concentration of approximately 330 µg/mL. BPs were synthesized by gradually adding acetone (Sigma-Aldrich) dropwise to the bone stock solution at a 3:1 acetone -to-solution ratio under constant stirring. After particle formation, BPs were washed with distilled water through four rounds of centrifugation, resuspended by ultrasonication, and freeze-dried at -80°C to obtain powder-form BPs for cell experiments.

For fluorescence-based monitoring of cellular uptake and intracellular retention, Texas Red-labeled BPs (Red-BPs) were synthesized by incorporating 5% Texas Red dye (ThermoFisher) into the bone stock solution before particle synthesis, following the same fabrication steps to maintain consistent properties.

### 2.2. Cell culture, IR treatment, and BP co-culture

BMSCs purchased from Lonza Bioscience were cultured in Mesenchymal Stem Cell Growth Medium (MSCGM, Lonza) under standard conditions (37°C, 5% CO_2_, and 95% humidity), with media changes every three days. The third passage BMSCs, following detachment with 0.25% trypsin-EDTA (Invitrogen), were seeded at a density of 1 × 10^5^ cells per well in 12-well plates and divided into three experimental groups:

Control group: BMSCs without IR exposure or BP co-culture.

IR group: BMSCs exposed to IR treatment.

IR&BP group: BMSCs exposed to IR followed by BP co-culture.

After 24 hours of seeding, BMSCs in the IR and IR&BP groups were exposed to a single 10 Gy dose of X-ray irradiation using a Precision CellRad X-ray source in Dr. Himburg’s Radiation Oncology laboratory. After IR exposure, cells in the IR group were maintained in standard MSCGM medium. In the IR&BP group, BPs were resuspended in MSCGM medium by sonication (final concentration: 20 μg/mL, based on BP dry weight) and added to the cells 4 hours post-IR. The BP-containing medium was maintained for 24 hours before being replaced with fresh MSCGM without BPs. To monitor BP uptake and retention, a subset of IR-treated BMSCs was treated with Red-BPs following the same protocol as the IR&BP group.

All groups were cultured under standard conditions (37°C, 5% CO_2_) for seven days post-IR, with media changes on days 1, 3, and 7.

### 2.3. Intracellular uptake and retention of BPs

After 1 and 7 days of co-culture, IR-treated BMSCs with Red-BPs were washed with PBS, fixed with 4% paraformaldehyde, and stained with CellMask™ Plasma Membrane Stain (Invitrogen) to visualize the cell membrane. Nuclei were counterstained with DAPI (1 µg/mL, Abcam) (n = 4 per group). Fluorescence imaging was performed using a Nikon A1 Laser Scanning Confocal Microscope (Keyence), and image analysis was carried out with Fiji software to assess BP uptake and intracellular retention over time.

### 2.4. Cell morphology and proliferation

Cell morphology in all groups was observed on days 1, 3, and 7 of culture using an inverted phase-contrast microscope (Keyence). To evaluate cell viability and proliferation, the Resazurin Assay Kit (Abcam) was employed. At each time point (n = 3 per group), the culture medium was replaced with fresh medium containing 44 μM resazurin and incubated at 37°C with 5% CO_2_ for 1 hour. Following incubation, 100 μL of the resazurin-containing medium was collected and transferred to black, opaque 96 -well plates (Corning) for fluorescence measurement, while fresh medium was added back to the wells to continue cell culture. Fluorescence intensity was measured using a monochromator-based spectrophotometer (Cytation 3, BioTek) with excitation at 540 nm and emission at 590 nm.

### 2.5. Alizarin Red S and Oil Red O staining

To evaluate osteogenic differentiation, BMSCs from all groups were cultured for 7 days and stained with Alizarin Red S (ARS) (ScienCell) (n = 4 per group). Cells were fixed with 4% paraformaldehyde (PFA) for 15 minutes at room temperature, washed with PBS, and incubated with 40 mM ARS solution for 30 minutes at room temperature. After staining, cells were washed five times with deionized water to remove excess dye.

For adipogenic differentiation assessment, BMSCs from all groups were similarly cultured for 7 days and stained using Oil Red O (Millipore) (n = 3 per group). Cells were fixed with 4% PFA for 15 minutes at room temperature, washed three times with PBS, and briefly rinsed with 50% ethanol for 20 seconds to remove residual water. Oil Red O staining was performed by incubating the cells for 15 minutes, followed by washing with 50% ethanol for 20 seconds and multiple PBS washes to remove excess dye.

### 2.6. Immunocytochemistry staining of cells

After 7 days of culture, BMSCs from all groups were prepared for immunocytochemistry. Cells were washed with PBS, fixed with 4% PFA for 15 minutes at room temperature, and permeabilized with 0.1% Triton X-100 in 0.1% sodium citrate on ice for 2 minutes. Apoptotic cells were labeled using the TUNEL assay (In Situ Cell Death Detection Kit, Roche) following the manufacturer’s instructions. All the cells were blocked with 3% bovine serum albumin (BSA, Sigma-Aldrich) for 1 hour at room temperature and incubated overnight at 4°C with primary antibodies against Ki67 (1:400, Abcam), γ-H2AX (1:200, ThermoFisher), and RAD51 (1:200, Abcam) diluted in blocking buffer. The next day, cells were washed with PBS and incubated for 1 hour at room temperature with appropriate secondary antibodies and phalloidin (1:400, Invitrogen) to stain F-actin. Nuclei were counterstained with DAPI (1 μg/mL, Sigma-Aldrich), followed by final PBS washes. Fluorescence images were acquired using a Nikon A1 laser scanning confocal microscope (Keyence) and analyzed with Fiji software.

### 2.7. RNA extraction, bulk RNA sequencing, and data analysis

After 7 days of culture, total RNA was extracted from all experimental groups (n = 3 per group) using the RNeasy Plus Mini Kit (Qiagen) following the manufacturer’s instructions. RNA concentration was measured using a Nanodrop ND-1000 spectrophotometer, and RNA quality was assessed by capillary electrophoresis (Agilent Technologies). RNA sequencing (RNA-Seq) was performed by Novogene Corporation Inc. Libraries were prepared using the KAPA Stranded RNA-Seq Library Prep Kit (Illumina), and sequencing was conducted on the NovaSeqXPlus platform (Illumina) to generate paired-end reads.

Adapter sequences were trimmed using TrimGalore (Version 0.4.4; parameters: --stringency 3, -q 20, paired-end mode). The quality of the trimmed reads was verified using FastQC (Version 0.11.5). Alignment of the bulk RNA-Seq data to the human reference genome (GRCh38) was performed using the STAR pipeline, which also quantified transcript abundances to account for variability due to read alignment [49, 50].

Differential gene expression (DEG) analysis was conducted using the R package DESeq2 (v1.42. 1) [51]. Statistical significance of expression differences between groups (IR vs Control and IR&BP vs IR) was determined using a likelihood ratio test. P-values were adjusted for multiple hypothesis testing using the Benjamini-Hochberg method to calculate the false discovery rate (FDR), and transcripts with FDR < 0.05 were considered significantly differentially expressed [52].

Visualization of differentially expressed transcripts was performed using the EnhancedVolcano R package (v1.20.0) to generate volcano plots, and the VennDiagram package was used to depict overlaps in DEG s and enriched pathways between group comparisons [53]. For DEG heatmaps, counts per million (CPM) estimates were calculated, and hierarchical clustering (complete linkage) was applied to visualize expression patterns among DE transcripts.

For functional analysis, over-representation analysis (ORA) was performed using the R package clusterProfiler (v4.10.1) to identify significantly enriched Gene Ontology (GO) terms related to biological processes. Enriched genes from overlapping upregulated and downregulated DEGs were selected based on the IR vs Control and IR&BP vs IR comparisons. The ComplexHeatmap package (v2.18.0) was used to generate annotated heatmaps based on Z-scores derived from CPM values. All plots were further customized and enhanced using the ggplot2 package (v3.5.1).

## 3. Results

### 2.1. Intracellular uptake and retention of BPs

The uptake and retention of Red-BPs in IR-treated BMSCs were assessed at days 1 and 7 using confocal fluorescence microscopy. After 1 day of co-culture with Red-BPs, bright red fluorescent signals were observed in the cytoplasm of BMSCs (Fig. 1A, left), confirming efficient internalization. Following this, the Red -BP- containing medium was replaced with fresh MSCGM without Red-BPs, and cells were maintained under standard culture conditions. By day 7, red fluorescence remained visible within the cytoplasm, although with slightly reduced intensity (Fig. 1A, right), indicating sustained intracellular retention of Red-BPs over time.

**Figure 1:**
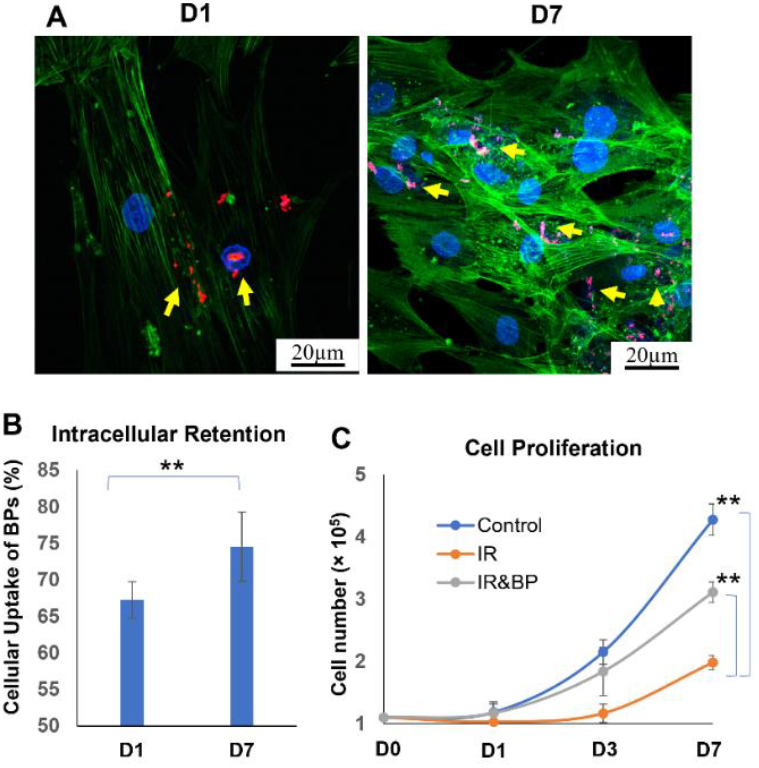
Intracellular Retention and Proliferation: (A) Confocal fluorescence microscopy images showing the uptake and intracellular retention of Red-BPs in IR-treated BMSCs at day 1 (left) and day 7 (right). (B) Quantification of Red-BP-positive cell numbers based on confocal fluorescence imaging. (C) Assessment of cell viability and proliferation using the Resazurin Assay. Statistical significance: *P < 0.05, **P < 0.01.

Quantitative analysis of confocal images (Fig. 1B) showed that 67.3 ± 2.5% of BMSCs had internalized Red-BPs by day 1. Interestingly, this proportion increased to 74.5 ± 4.7% by day 7, suggesting not only retention but also potential redistribution of Red-BPs as the cells proliferated.

### 3.2. Cell morphology and proliferation

The effects of IR exposure and BP co-culture on BMSC morphology and proliferation were evaluated using phase contrast microscopy and the Resazurin Assay. In the control group, BMSCs maintained a typical spindle-shaped morphology with uniform distribution and progressive increases in cell density over 7 days, indicating healthy growth and viability (Fig. 2A).

**Figure 2:**
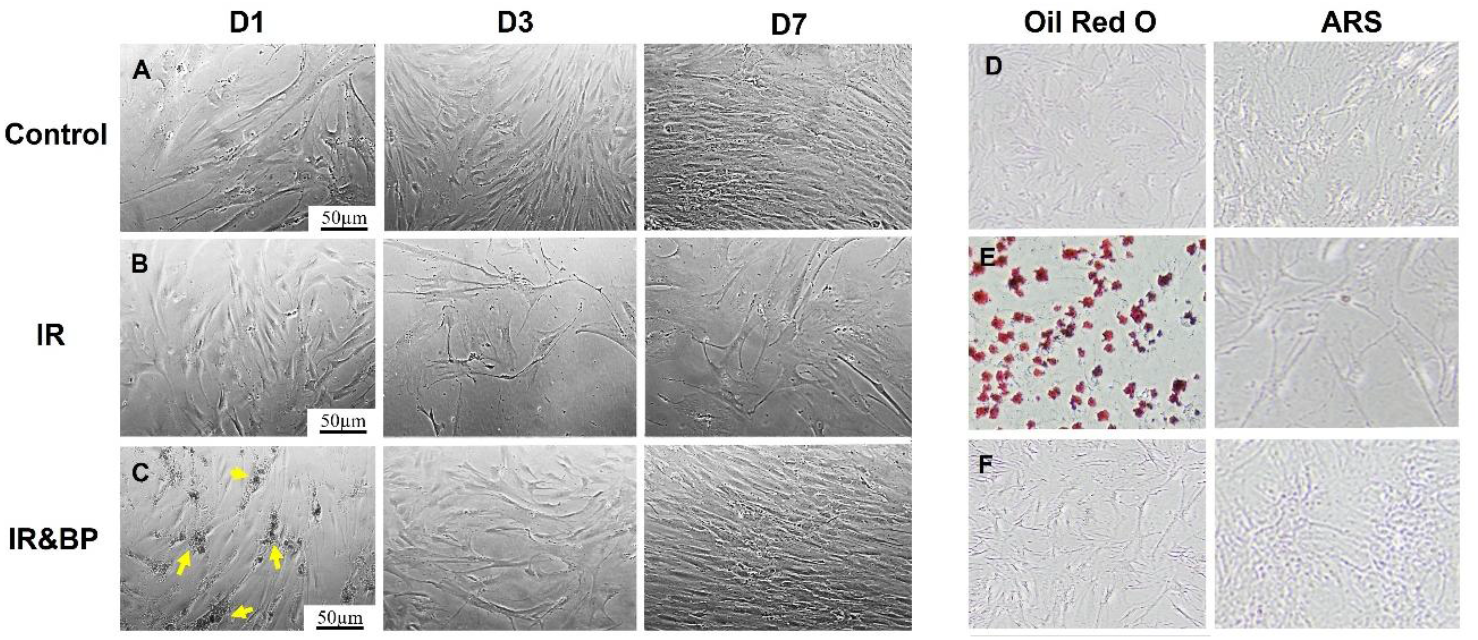
Morphology and Spontaneous Differentiation Potential of BMSCs: Phase-contrast microscopy images showing BMSC morphology in the (A) control group, (B) IR group, and (C) IR&BP group. Evaluation of spontaneous differentiation potential after 7 days of culture without specific induction. Cells were stained with Oil Red O (left panels) for adipogenic differentiation and Alizarin Red S (ARS) (right panels) for mineralization in the (D) control group, (E) IR group, and (F) IR&BP group.

In contrast, BMSCs in the IR group showed a thinner, more elongated morphology and a marked reduction in cell density at both day 3 and day 7 post-IR exposure (Fig. 2B), consistent with IR-induced damage and impaired growth.

In the IR&BP group (Fig. 2C), BPs were visibly internalized by BMSCs by day 1. By day 3, cell density increased compared to the IR group, and by day 7, BMSCs regained their typical spindle -shaped morphology and formed a well-organized monolayer, closely resembling the control group, indicating that BP treatment supported cell survival and partially restored proliferation following IR-induced damage.

Cell viability and proliferation were further assessed using the Resazurin Assay (Fig. 1C). The IR group exhibited significantly reduced growth rates and lower cell numbers compared to the control group. Although the IR&BP group did not fully restore proliferation to control levels, it showed substantial improvement in cell viability and growth compared to the IR group after 7 days. Together, these results demonstrate that BP co-culture mitigates IR-induced cellular damage by promoting BMSC survival and enhancing proliferation.

### 3.3. Potential osteogenic and adipogenic differentiation of BMSCs

To evaluate the effects of IR exposure and BP treatment on the spontaneous differentiation potential of BMSCs, cells were stained with Oil Red O (for adipogenic differentiation) and ARS (for osteogenic differentiation) after 7 days of culture without specific induction. Oil Red O staining revealed minimal lipid droplet formation in both the control group (Fig. 2D, left) and the IR&BP group (Fig. 2F, left), indicating a low level of adipogenic differentiation. In contrast, the IR group (Fig. 2E, left) exhibited a significantly greater number of Oil Red O-positive cells, suggesting that IR exposure promoted adipogenic differentiation. ARS staining showed no notable mineralized nodule formation in any of the groups (Fig. 2D–F, right), indicating that osteogenic differentiation was limited under the current culture conditions across all groups.

### 3.4. Evaluation of cell proliferation and apoptosis

To assess the effects of IR exposure and BP treatment on BMSC proliferation and apoptosis, cells were stained for Ki67 (a proliferation marker) and TUNEL (an apoptosis marker) after 7 days of culture. In the control group, the majority of BMSCs were strongly positive for Ki67, with very few TUNEL-positive cells detected (Fig. 3A), indicating robust proliferation and minimal apoptosis. In contrast, most of the cells in the IR group exhibited TUNEL-positive staining (Fig. 3B) with a significant increase in cells positive for TUNEL (Fig. 3H, P<0.0001) along with markedly reduced Ki67 expression (Fig. 3G, P<0.0001) compared to the control group, consistent with extensive IR-induced apoptosis and impaired proliferative activity. In the IR&BP group, although some TUNEL-positive cells (Fig. 3C, arrow) were still present, their number was significantly lower compared to the IR group (Fig. 3H, P<0.0001) and not statistically different from the control group (Fig. 3H, P=0.17). Additionally, Ki67 expression in the IR&BP group (Fig. 3C) was comparable to that of the control group (Fig. 3G, P=0.77), suggesting that BP treatment effectively mitigated IR-induced apoptosis and preserved the proliferative capacity of BMSCs.

**Figure 3:**
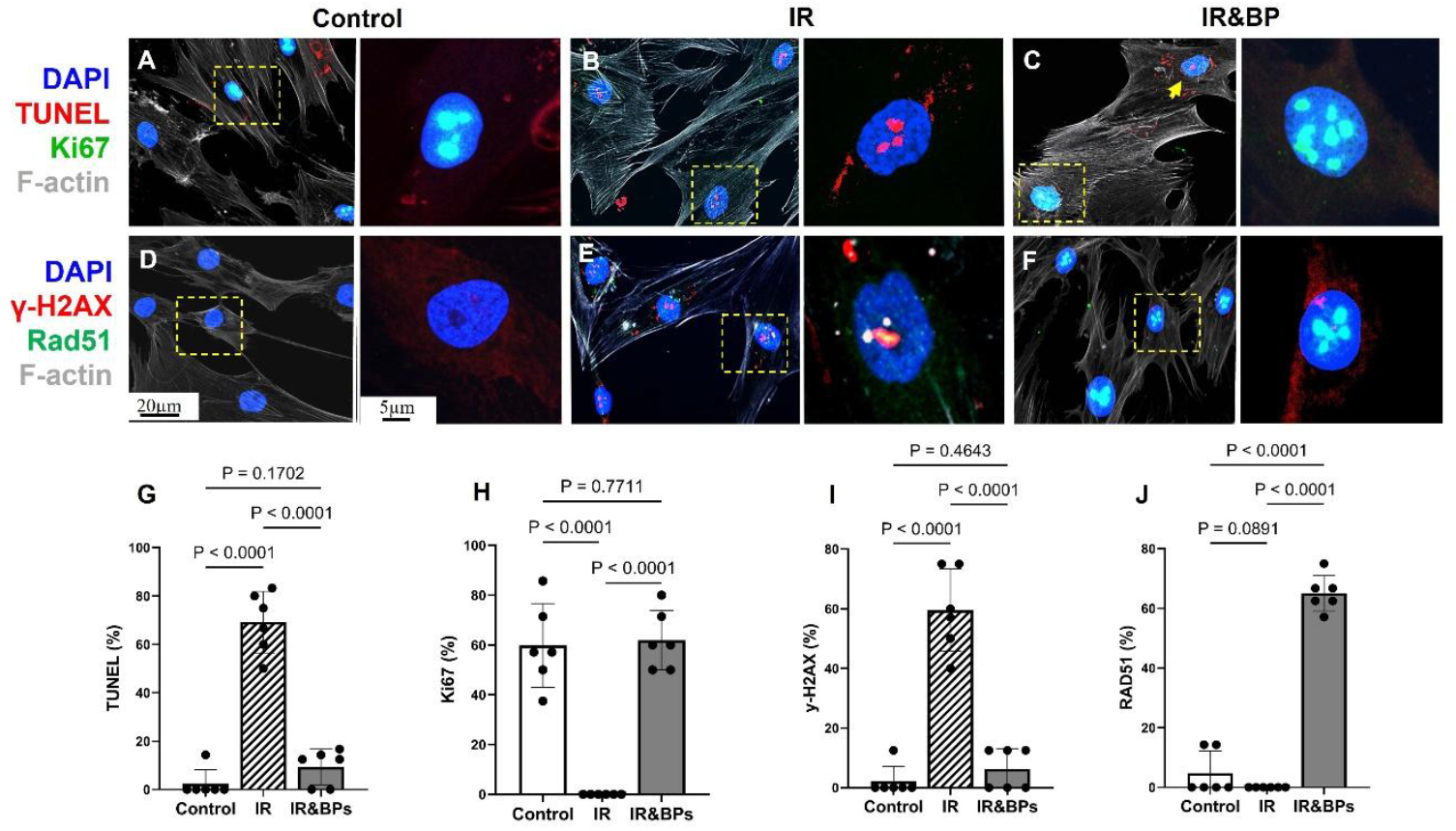
Evaluation of BMSC Proliferation, Apoptosis, DNA Damage, and DNA Repair: Immunofluorescence staining to assess proliferation and apoptosis after 7 days of culture in the (A) control group, (B) IR group, and (C) IR&BP group. Cells were stained for Ki67 (green, proliferation marker), TUNEL (red, apoptosis marker), F-actin (gray, cytoskeleton), and DAPI (blue, nuclei). Immunofluorescence staining to assess DNA damage and repair after 7 days of culture in the (D) control group, (E) IR group, and (F) IR&BP group. Cells were stained for γ-H2AX (red, DNA double-strand break marker), Rad51 (green, DNA repair marker), F-actin (gray), and DAPI (blue). Quantitative analysis of the percentage of positive cells for (G) TUNEL, (H) Ki67, (I) γ- H2AX, and (J) Rad51.

### 3.5. Evaluation of DNA damage and repair

To further investigate the effects of IR exposure and BP treatment on DNA damage and repair in BMSCs, cells were stained for γ-H2AX (a marker of DNA double-strand breaks) and Rad51 (a marker of DNA repair activity) after 7 days of culture. In the control group, very few γ-H2AX-positive foci were observed, indicating minimal baseline DNA damage (Fig. 3D). Correspondingly, Rad51 expression was also low, consistent with the low requirement for DNA repair under normal conditions. Following IR exposure, BMSCs in the IR group showed a substantial increase in γ-H2AX foci formation (Fig. 3E), with a significantly higher percentage of γ-H2AX- positive cells compared to the control group (Fig. 3I, P<0.0001), reflecting extensive DNA damage. Although Rad51 expression was detectable in the IR group, the number of Rad51-positive cells remained low, suggesting a limited DNA repair response following IR-induced damage. In contrast, BMSCs in the IR&BP group exhibited a marked reduction in γ-H2AX-positive staining (Fig. 3F), which was significantly lower than in the IR group (Fig. 3I, P<0.0001), indicating that BP treatment alleviated persistent DNA damage. Moreover, Rad51 expression in the IR&BP group was significantly enhanced compared to the IR group (Fig. 3J), suggesting th at BP treatment promoted the activation of DNA repair.

### 3.6. Transcriptome changes in BMSCs

RNA sequencing analysis demonstrated distinct transcriptomic alterations among BMSCs from the control, IR, and IR&BP groups (Fig. 4A). Using a false discovery rate (FDR)-corrected p-value of 0.05 and a fold change (FC) threshold of 1.5, we identified significant changes in gene expression profiles. Compared to the control group, BMSCs in the IR group exhibited 1,192 upregulated and 871 downregulated genes (Fig. 4B), reflecting the extensive impact of IR exposure. When comparing the IR&BP group to the IR group, a more pronounced transcriptomic shift was observed, with 4,090 genes upregulated and 4,082 genes downregulated (Fig. 4C), suggesting that BP treatment substantially altered gene expression following IR injury.

**Figure 4:**
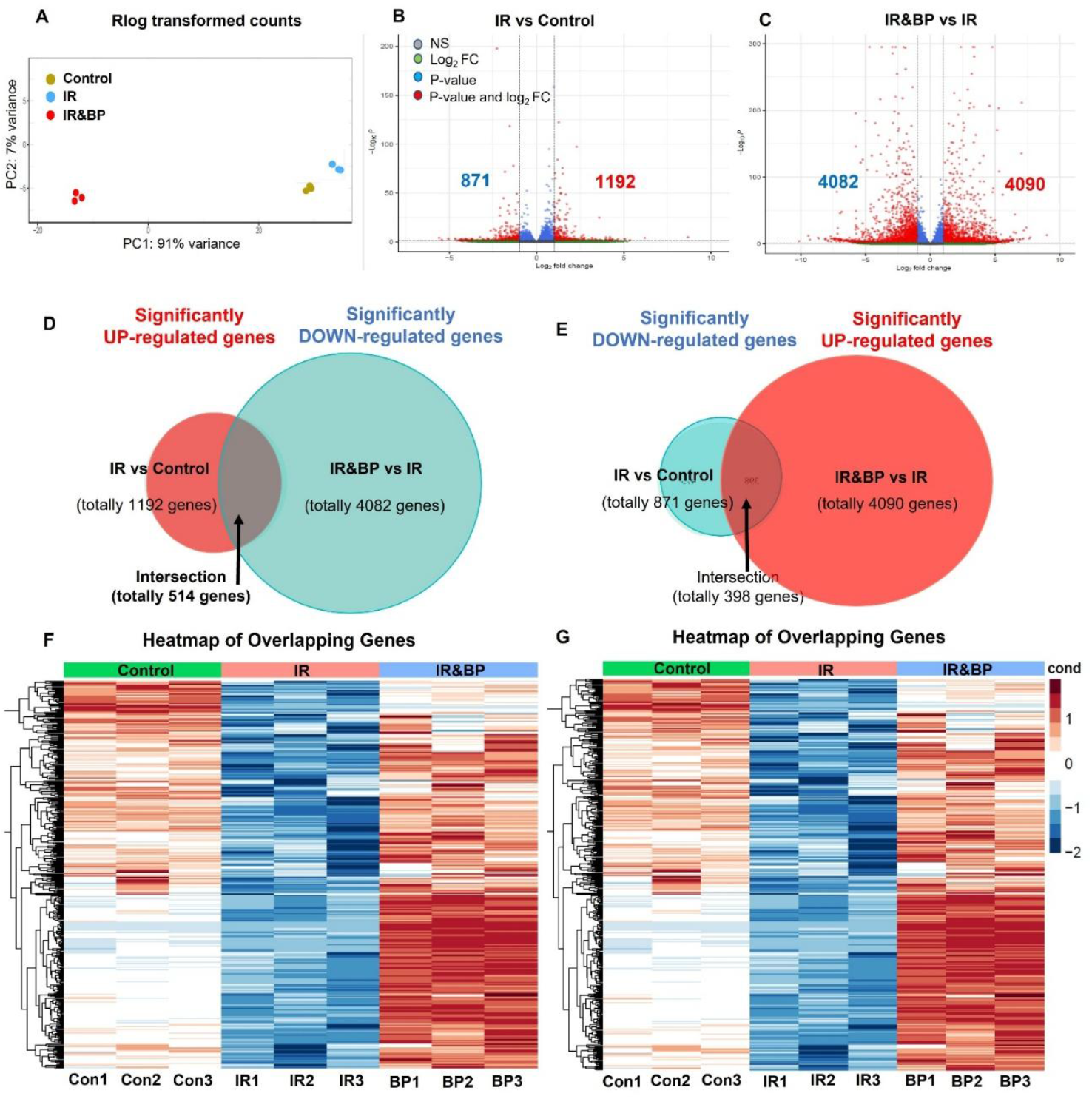
Differential Gene Expression Analysis in IR and IR&BP Groups: (A) Principal component analysis (PCA) showing the transcriptomic separation among the control, IR, and IR&BP groups. Volcano plots of differentially expressed genes (DEGs) of (B) IR group vs. control group and (C) IR&BP group vs. IR group. The horizontal line represents the false discovery rate (FDR)-corrected p-value threshold of 0.05, and the vertical lines indicate fold change (FC) cutoffs of 1.5. Venn diagrams illustrating overlapping DEGs of (D) Genes significantly upregulated by IR and downregulated by BP treatment and (E) Genes significantly downregulated by IR and upregulated by BP treatment. Heatmaps showing the expression patterns of intersection genes of (F) 514 genes from panel D and (G) 398 genes from panel E.

Further intersection analysis identified 514 genes that were significantly upregulated in the IR group but downregulated following BP treatment (Fig. 4D), as illustrated in the heat map (Fig. 4F). Conversely, 398 genes were significantly downregulated in the IR group but upregulated in the IR&BP group (Fig. 4E), with corresponding heat maps shown in Fig. 4G. These results indicate that BP treatment not only mitigates IR - induced transcriptomic changes but may also activate specific gene pathways related to recovery and repair.

### 3.7. Functional enrichment analysis

To further evaluate the functional consequences of IR exposure and BP treatment on BMSCs, we performed pathway enrichment analysis focusing on the top 10 most significantly upregulated and downregulated pathways for each group.

In the IR group, the upregulated pathways (Fig. 5A) were primarily associated with extracellular matrix (ECM) metabolism and remodeling, including ECM organization, glycoprotein metabolism and biosynthesis, collagen fibril organization, glycosaminoglycan metabolism, proteoglycan metabolism, mucopolysaccharide metabolism, and aminoglycan metabolism. These results suggest that IR exposure induces enhanced ECM remodeling, likely as a compensatory response to IR-induced cellular stress. The downregulated pathways in the IR group (Fig. 5B) were largely related to cell cycle regulation, including mitotic sister chromatid segregation, mitotic nuclear division, chromosome segregation, and chromosome organization, indicating that IR exposure impairs normal mitotic processes and disrupts proliferative capacity.

**Figure 5:**
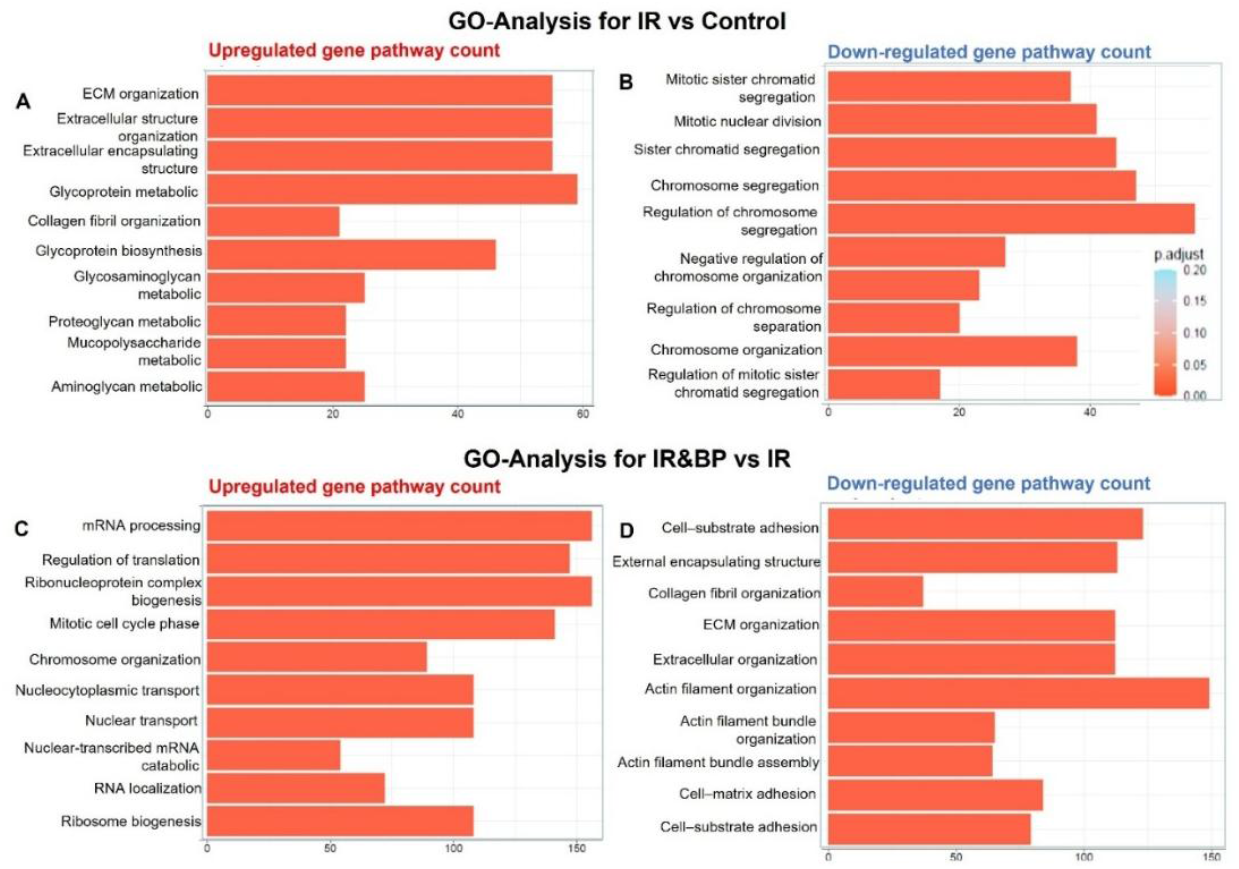
Gene Ontology (GO) Enrichment Analysis of DEGs in IR and IR&BP Groups: (A) Top 10 significantly upregulated pathways identified in the IR group based on GO enrichment analysis. (B) Top 10 significantly downregulated pathways in the IR group. (C) Top 10 significantly upregulated pathways in the IR&BP group. (D) Top 10 significantly downregulated pathways in the IR&BP group.

In contrast, the IR&BP group exhibited adistinct transcriptomic shift. The top 10 upregulated pathways (Fig. 5C) were primarily associated with RNA processing and cell cycle recovery, including mRNA processing, translation regulation, ribonucleoprotein complex biogenesis, mitotic cell cycle phase transition, chromosome organization, nucleocytoplasmic and nuclear transport, RNA localization, and ribosome biogenesis. These findings suggest that BP treatment not only mitigates IR-induced cell cycle arrest but actively promotes the restoration of proliferative and biosynthetic functions. Additionally, the downregulated pathways in the IR&BP group were mainly related to cell-substrate adhesion, ECM organization, collagen fibril organization, actin filament organization, and cell-matrix adhesion (Fig. 5D), indicating that BP treatment suppresses the IR-induced ECM remodeling response.

To investigate the shift in gene expression between the IR and IR&BP groups, we conducted a comparative pathway enrichment analysis. Comparison of the IR group versus Control revealed a total of 163 significantly upregulated pathways, while IR&BP versus IR exhibited 211 significantly downregulated pathways (Fig. 6A). Notably, 35 pathways overlapped between the two comparisons, suggesting that IR&BP treatment specifically counteracts many of the pathways activated by IR treatment.

**Figure 6.**
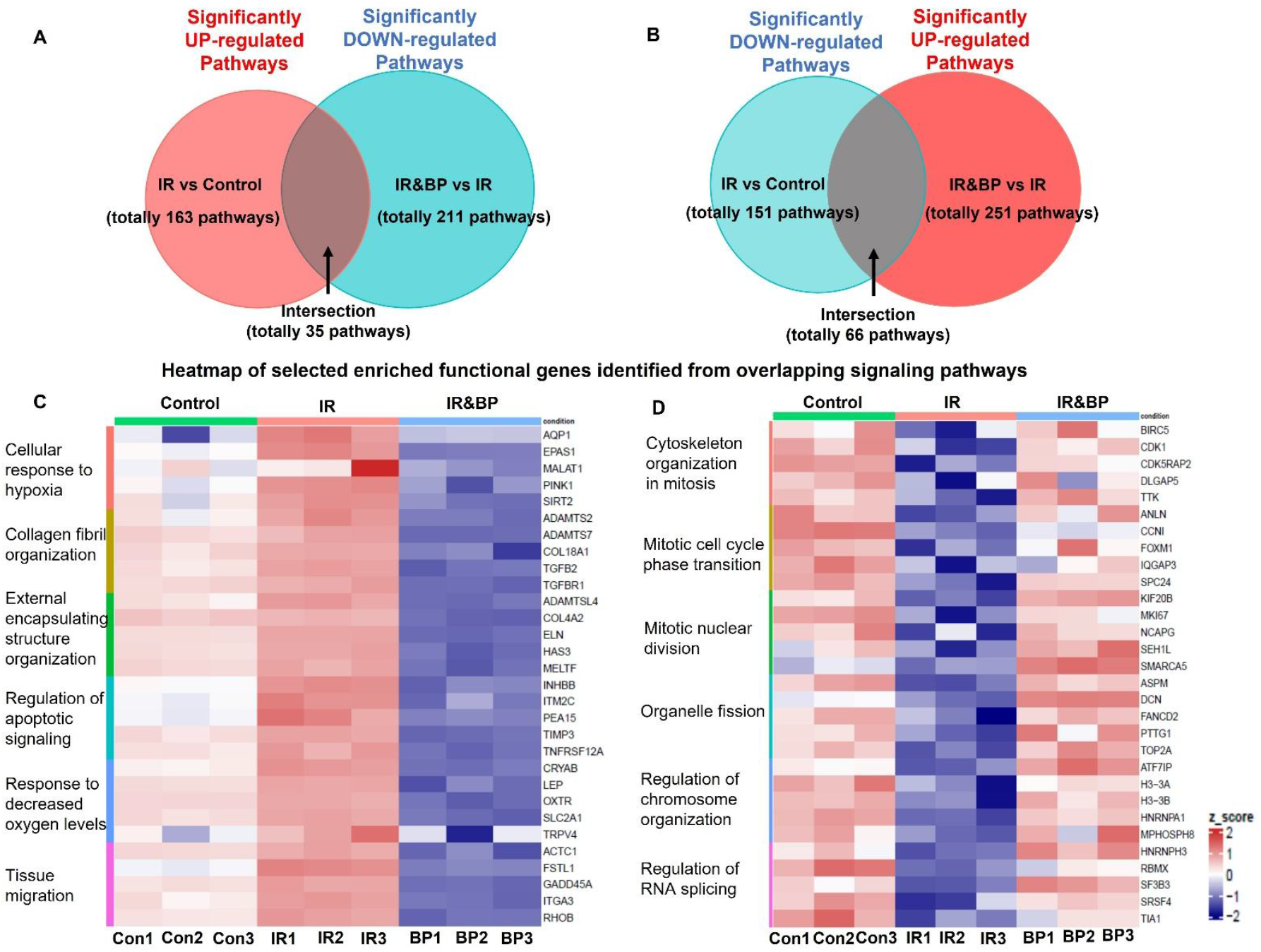
Comparative Pathway Enrichment and Gene Expression Analysis: (A) Venn diagram showing the overlap between pathways significantly upregulated in the IR group and significantly downregulated in the IR&BP group. (B) Venn diagram showing the overlap between pathways significantly downregulated in the IR group and significantly upregulated in the IR&BP group. (C) Heatmap of representative genes from overlapping pathways that were upregulated by IR but downregulated by BP treatment. (D) Heatmap of representative genes from overlapping pathways that were downregulated by IR but upregulated by BP treatment.

To further explore the functional significance of these pathway alterations, we generated a heatmap depicting the expression profiles of selected enriched genes from the overlapping pathways (Fig. 6C). IR markedly upregulated genes involved in cellular stress responses, ECM remodeling, and tissue organization. Notably, *EPAS1* (hypoxia-inducible factor 2α), *SIRT2* (a stress-response regulator), and several ECM-related genes, including *COL18A1, TGFβ2, ELN*, and *HAS3*, were significantly elevated following IR exposure. In contrast, IR&BP group dramatically downregulated the expression of these genes, bringing their levels back toward or even below those observed in the control group. For example, genes associated with collagen fibril organization (*COL18A1, COL4A2*), external encapsulating structure (*ELN, HAS3*), and response to hypoxia (*EPAS1, PINK1*) were substantially normalized following IR&BP treatment. Furthermore, key regulators of apoptotic signaling (*TIMP3, TNFRSF12A*) and tissue migration (*GADD45A, ITGA3, RHOB*) also showed a reversal of IR-induced expression changes following BP treatment.

Conversely, IR versus Control also resulted in significant downregulation of 151 pathways, whereas IR&BP versus IR comparison revealed significant upregulation of 251 pathways, with 66 pathways overlapping between the two groups (Fig. 6B).

To investigate the functional relevance of the rescued pathways, we generated a second heatmap focusing on key genes from these overlapping pathways (Fig. 6D). IR exposure led to a marked downregulation of genes critical for mitotic progression and cell division, including *BIRC5* (survivin), *CDK1* (cyclin-dependent kinase 1), *ANLN* (anillin), and *FOXM1* (a master regulator of cell cycle progression). In addition, genes involved in mitotic spindle formation and chromosome segregation, such as *DLGAP5, KIF20B, SPC24*, and *NCAPG*, were also significantly suppressed by IR treatment. In the IR&BP group, the expression of these genes was substantially restored, approaching or even exceeding levels observed in the control group. Moreover, genes associated with DNA replication and repair (*TOP2A, FANCD2, PTTG1*) and RNA processing and splicing (*HNRNPA1, SF3B3, TIA1*) were significantly upregulated in the IR&BP group, further supporting the notion that BP treatment promotes cellular recovery mechanisms after IR-induced damage.

## 4. Discussion

IR-induced bone damage represents a significant clinical challenge, particularly for patients undergoing radiotherapy and individuals exposed to high doses of radiation [17-20]. A major contributor to IR-induced bone damage is its detrimental effect on BMSCs, which impairs their osteogenic differentiation capacity and increases apoptosis, ultimately leading to bone fragility, decreased mineral density, structural degradation, and a heightened risk of fractures [35]. Given these challenges, there is a pressing need for therapeutic strategies that can effectively reverse IR-induced BMSC dysfunction. Specifically, approaches that restore the proliferative capacity and osteogenic potential of BMSCs are essential to support bone regeneration following IR exposure. In this study, we investigated the effects of IR on BMSC function and evaluated BPs as a potential nanotherapeutic intervention to mitigate IR-induced cellular damage and promote bone repair.

IR is well established to cause DNA damage, induce apoptosis, and trigger cell cycle arrest across a variety of cell types [54-56]. Preclinical models commonly apply X-ray doses ranging from 2 to 10 Gy to study these effects [57-60]. In this study, we selected a 10 Gy dose to model severe IR-induced cellular injury, providing a stringent and physiologically relevant platform for evaluating BMSC responses. To systematically assess both the extent of IR-induced cellular damage and the potential protective effects of BPs, BMSCs were divided into three experimental groups: a control group (untreated), an IR group (exposed to 10 Gy X-ray), and an IR&BP group (exposed to 10 Gy IR followed by intracellular delivery of BPs).

As expected, BMSCs in the IR group exhibited pronounced morphological and functional impairments compared to the control group. IR-treated cells appeared thinner, more elongated, and less spread out, suggesting cytoskeletal disruption. Cell proliferation was also significantly reduced along with a decrease in the number of Ki67-positive cells, indicating impaired cell proliferation. IR exposure also led to extensive DNA damage, demonstrated by a substantial increase in γ-H2AX foci formation and a significantly higher proportion of γ-H2AX-positive cells compared to controls. Additionally, an increased number of TUNEL-positive cells indicated elevated levels of apoptosis. Collectively, these findings confirm that 10 Gy IR induces profound damage to BMSCs, severely impairing their proliferative and regenerative capacity.

To evaluate the therapeutic potential of BPs in mitigating IR-induced cellular damage, BPs were administered to BMSCs in the IR&BP group at a concentration of 20 μg/mL, 4-hours after 10 Gy IR exposure. After a 24-hour incubation period to allow intracellular uptake, the medium was replaced with fresh culture media. To monitor BP internalization and retention, Red-BPs were used and visualized via confocal microscopy. By day 1, strong red fluorescence localized to the cytoplasm confirmed efficient intracellular uptake of BPs. Notably, the fluorescence signal persisted up to day 7, although with slightly reduced intensity, suggesting sustained intracellular retention. Furthermore, the increased proportion of Red-BP-positive cells over time implies potential redistribution or inheritance of BPs during cell proliferation. These findings support the effective delivery and long-term intracellular presence of BPs, providing a basis for their therapeutic activity in protecting BMSCs against IR-induced damage.

Functionally, BP treatment significantly improved the morphology, viability, and proliferation of BMSCs after IR exposure, as evidenced by a higher rate of cell proliferation, increased Ki67-positive cells, and a reduction in TUNEL-positive cells, indicating both enhanced proliferation and reduced apoptosis. Moreover, BP treatment effectively alleviated persistent DNA damage, as shown by a marked reduction in γ-H2AX-positive staining and increased Rad51 expression in the IR&BP group compared to the IR-only group. These findings suggest that BP treatment enhances DNA repair through homologous recombination, mitigating the DNA damage caused by IR exposure [61, 62]. Collectively, these results indicate that internalized BPs help counteract IR-induced cellular stress by improving cell proliferation and facilitating DNA repair, thereby promoting the recovery of BMSCs after IR injury.

To evaluate the impact of IR exposure and BP treatment on the spontaneous differentiation potential of BMSCs, Oil Red O and ARS staining were performed after 7 days of IR exposure and cell culture without specific differentiation induction. Oil Red O staining revealed increased lipid droplet accumulation in the IR group, indicating a shift toward adipogenic differentiation. In contrast, the control and IR&BP groups showed minimal lipid accumulation, suggesting that BP treatment may counteract the IR-induced shift toward adipogenesis. ARS staining showed no significant mineralized matrix formation in any of the groups, indicating that spontaneous osteogenic differentiation did not occur under these conditions. These findings suggest that IR exposure promotes an undesirable shift in BMSC differentiation toward adipocytes. However, BP co -culture appears to help preserve a more balanced differentiation potential, supporting the maintenance of osteogenic capacity for future regenerative applications.

In summary, the above findings showed that BMSCs exposed to high doses of IR undergo significant morphological and functional impairments, including reduced proliferation, increased apoptosis, persistent DNA damage, and a shift toward adipogenic differentiation. However, BP treatment to BMSCs effectively mitigates these IR-induced damages by promoting cell proliferation, reducing apoptosis, alleviating DNA damage, and enhancing DNA repair. Additionally, BPs counteracted the IR-induced shift toward adipogenesis, helping to preserve the cells’ osteogenic potential. This preservation is crucial for bone regeneration, as excessive adipogenesis can impair skeletal repair, particularly following IR exposure or with aging.

BPs represent a novel class of protein-based nanoparticles derived from decellularized bone, a material widely used in orthopedic applications [63-67]. The decellularization process effectively removes immunogenic cellular components while preserving essential ECM proteins, such as collagen and osteoinductive growth factors, which are critical for bone repair and regeneration [68-71]. Our previous studies have demonstrated that BPs retain the biocompatibility and biodegradability of bone while offering a nanoscale structure that enhances cellular uptake and interaction [46-48]. A major advantage of BPs is their ability to mimic the natural ECM environment, which is critical for regulating stem cell functions, including survival, migration, and differentiation. By replicating these biological cues at the nanoscale, BPs provide intracellular support once internalized by BMSCs. This support is especially beneficial under stress conditions, such as those induced by IR, as BPs help reduce DNA damage, activate pro-survival pathways, and maintain balanced differentiation. These protective effects make BPs a promising therapeutic tool for enhancing stem cell function and promoting bone regeneration, particularly in settings of IR-induced injury where preserving the osteogenic potential of BMSCs is vital for successful skeletal repair.

Given the promising protective effects of BPs on BMSCs under IR-induced stress, we further investigated the underlying molecular mechanisms using RNA sequencing to examine transcriptional changes and pathway alterations associated with IR-induced dysfunction and BP-mediated recovery.

RNA sequencing analysis revealed that IR exposure caused substantial transcriptomic disruptions in BMSCs, characterized by upregulation of ECM remodeling pathways and downregulation of genes involved in cell cycle regulation and mitosis, suggesting impaired proliferative capacity and activation of stress responses. This transcriptomic shift aligns with the previously observed morphological and functional impairments—reduced proliferation, increased apoptosis, and loss of differentiation balance —indicating that IR not only damages BMSCs at the cellular level but also fundamentally alters their molecular programs, compromising their regenerative potential.

BP treatment markedly altered this trajectory, reversing many IR-induced transcriptomic changes. BPs suppressed pathways associated with ECM remodeling and hypoxia response while restoring pathways critical for cell cycle progression, DNA repair, and RNA processing. Key genes regulating mitosis and proliferation, such as CDK1, FOXM1, and BIRC5, were significantly upregulated following BP treatment, suggesting improved cell division and recovery. Notably, BPs normalized the expression of stress- and apoptosis-related genes (e.g., EPAS1, TIMP3) while enhancing genes involved in DNA replication and chromosomal stability (e.g., TOP2A, FANCD2). These results indicate that BPs promote a shift from a stress -induced, dysfunctional state to a regenerative, proliferative phenotype. Notably, BPs normalized the expression of stress - and apoptosis-related genes (e.g., EPAS1, TIMP3) while enhancing genes involved in DNA replication and chromosomal stability (e.g., TOP2A, FANCD2). Together, these findings demonstrate that BPs not only protect BMSCs from IR-induced damage but actively restore their proliferation and regenerative potential, highlighting their therapeutic promise in IR-related bone repair applications.

While this study provides important insights into the protective effects of BPs on BMSCs under IR-induced stress, several limitations and areas for future research remain. Although RNA sequencing revealed significant transcriptional changes following IR exposure and BP treatment, it remains unclear how these gene expression alterations translate into functional outcomes. Future studies incorporating proteomic and metabolomic analyses would help clarify how BP treatment influences key cellular processes such as protein synthesis, energy metabolism, and ECM remodeling. This would provide a more comprehensive understanding of the molecular mechanisms driving BP-mediated protection and recovery. Furthermore, while BP treatment appeared to reverse many IR-induced transcriptomic disruptions, the long-term effects on BMSC functionality—particularly their capacity for osteogenic differentiation and integration into bone tissue —require further investigation to fully assess the therapeutic potential of BPs in regenerative medicine.

## 5. Conclusion

In conclusion, this study highlighted the potential of BPs as a novel nanotherapeutic strategy to protect and restore BMSCs following IR exposure. BMSCs exposed to high doses of IR showed significant impairments, including reduced proliferation, increased apoptosis, persistent DNA damage, and a shift toward adipogenic differentiation, all of which compromised their regenerative capacity. BP treatment effectively mitigated these effects by promoting proliferation, reducing apoptosis, alleviating DNA damage, enhancing DNA repair, and preserving osteogenic differentiation potential.

At the molecular level, RNA sequencing analysis revealed that BPs reversed critical transcriptomic alterations induced by IR, notably restoring pathways involved in cell cycle progression, ECM remodeling, and apoptotic regulation. By re-establishing genomic stability and functional integrity, BPs significantly supported the recovery and regenerative potential of BMSCs after IR injury.

These findings position BPs as a promising platform for enhancing stem cell-based bone regeneration, particularly in clinical scenarios involving IR-induced skeletal damage. Their natural ECM-mimetic composition, biocompatibility, and nanoscale delivery properties offer distinct advantages for regenerati ve medicine applications. Future research should focus on elucidating the detailed molecular mechanisms of BP-mediated protection, validating efficacy in *in vivo* models of bone repair, and investigating combinatorial strategies with other regenerative therapeutics to advance clinical translation.

## Consent for publication

Not applicable.

## Availability of data and materials

The datasets generated and/or analyzed during the current study are available from the corresponding author on reasonable request.

## Funding and acknowledgements

This research was supported by a Startup Fund and a Product Development Pilot Award from the Joint Department of Biomedical Engineering at Marquette University and the Medical College of Wisconsin, awarded to Dr. Bo Wang.

## Author contributions

Conceptualization: BW

Methodology: BW, XM, CN, AS, SL, LH, AF, HH, LG, PD, SW, MM, RW

Investigation: BW, XM, CN, AS, SL, LH, AF, HH, LG, PD, SW, MM, RW

Visualization: BW

Supervision: BW

Writing: BW

## Data availability statement

The datasets generated and analyzed during the current study are available from the corresponding author upon reasonable request.

